# Acylcarnitine Metabolomic Profiles Inform Clinically-Defined Major Depressive Phenotypes

**DOI:** 10.1101/632448

**Authors:** Ahmed T. Ahmed, Siamak MahmoudianDehkordi, Sudeepa Bhattacharyya, Matthias Arnold, Duan Liu, Drew Neavin, M. Arthur Moseley, J. Will Thompson, Lisa St John Williams, Gregory Louie, Michelle K. Skime, Liewei Wang, Patricio Riva-Posse, William McDonald, William V. Bobo, W. Edward Craighead, Ranga Krishnan, Richard M. Weinshilboum, Boadie W. Dunlop, David S. Millington, A. John Rush, Mark A. Frye, The Mood Disorders Precision Medicine Consortium (MDPMC), Rima Kaddurah-Daouk

## Abstract

**Background:** Acylcarnitines have important functions in mitochondrial energetics and β-oxidation, and have been implicated to play a significant role in metabolic functions of the brain. This retrospective study examined whether plasma acylcarnitine profiles can help biochemically distinguish the three phenotypic subtypes of major depressive disorder (MDD)—(core depression (CD+), anxious depression (ANX+), and neurovegetative symptoms of melancholia (NVSM+))—following treatment with a selective serotonin reuptake inhibitor (SSRI).

**Methods:** Depressed outpatients (n=240) from the Mayo Clinic Pharmacogenomics Research Network were treated with citalopram or escitalopram for eight weeks. Plasma samples collected at baseline and eight weeks post-treatment were profiled for multiple-, short-, medium- and long-chain acylcarnitine levels using AbsoluteIDQ^®^p180-Kit and LC-MS. Linear mixed effects models were used to examine whether acylcarnitine levels discriminate the clinical phenotypes at baseline or eight weeks post-treatment, and whether temporal changes in acylcarnitine profiles differ between groups.

**Results:** At baseline, significantly lower concentrations of short- and long-chain acylcarnitines were found in CD+ and NVSM+ compared to ANX+, and the short-chain acylcarnitines remained lower after eight weeks. At eight weeks, the medium- and long-chain acylcarnitines were significantly lower in NVSM+ compared to ANX+. Regarding changes baseline to week eight, short-chain acylcarnitine levels significantly increased in CD+ and ANX+, and medium- and long-chain acylcarnitines significantly decreased in NVSM+ and CD+.

**Conclusions:** In depressed patients treated with SSRIs, β-oxidation and mitochondrial energetics as evaluated by levels and changes in acylcarnitines may provide the biochemical basis of the clinical heterogeneity of MDD, especially when combined with clinical characteristics.

## Introduction

Treatment response in patients with major depressive disorder (MDD) is only partially predicted by clinical symptom profiles, and depressive symptoms alone fail to account for the majority of biological heterogeneity in treatment response (Carragher, Adamson, Bunting, & McCann, 2009; Cassano et al., 2009; Charney & Drevets, 2002; Harald & Gordon, 2012). Recent efforts to better parse this heterogeneity have included molecular, physiological, imaging, and neuropsychological measures combined with clinical data, with the aim of facilitating more personalized patient care (A. John Rush & Hicham M. Ibrahim, 2018; Rush & Ibrahim, 2018). To support biomarker identification in MDD, we recently proposed three depressive phenotypes based on items from the 17-item Hamilton Rating Scale for Depression (HRSD_17_) (Ahmed et al., 2018) that may reflect distinct neural processes aligned with the Research Domain Criteria (RDoC) proposed by the National Institute of Mental Health (Insel et al., 2010). These phenotypes, with corresponding question numbers on the HRSD_17_, were categorized as: 1) core depressive (CD+: depressed mood #1, work and activities #7); 2) neurovegetative symptoms of melancholia (NVSM+: late insomnia #6, somatic symptoms gastrointestinal #12); and 3) anxious features (ANX+: agitation #9, psychological anxiety #10, somatic anxiety #11, and hypochondriasis #15).

The present study was undertaken to determine whether levels of acylcarnitines at baseline and after eight weeks of treatment with a selective serotonin reuptake inhibitor (SSRI), or changes in the acylcarnitine profiles from baseline to week eight would discriminate these phenotypes from the others. This report focuses on acylcarnitines due to their emerging role in the central nervous system and relevance to MDD (Chen et al., 2014). Acylcarnitines are involved in mitochondrial function and energy metabolism, anti-oxidative functions and membrane stability, gene expression, neurotransmission, and neuroprotection (Calabrese, Stella, Calvani, & Butterfield, 2006; Nałęcz, Miecz, Berezowski, & Cecchelli, 2004; Pettegrew, Levine, & McClure, 2000; Virmani & Binienda, 2004; ZANELLI, SOLENSKI, ROSENTHAL, & FISKUM, 2005). Carnitine is present in the brain (Bresolin, Freddo, Vergani, & Angelini, 1982; Shug, Schmidt, Golden, & Fariello, 1982), where it transfers acetyl groups formed during β-oxidation to the cytosol, and it assists with the transfer of fatty acids from cytosol to mitochondria for energy production (FRITZ & MCEWEN, 1959; Jones, McDonald, & Borum, 2010).

Elevated medium- and long-chain acylcarnitine concentrations in blood have been associated with incomplete β-oxidation of fatty acids in a rat model of depression (Chen et al., 2014). In clinical studies, patients with a lifetime diagnosis of MDD demonstrated altered mitochondrial function or reduced ATP production (Beasley et al., 2006; Gardner et al., 2003; Suomalainen et al., 1992). Further, patients with mitochondrial disorders frequently have depressive symptoms or a history of MDD (Fattal, Link, Quinn, Cohen, & Franco, 2007; Kato, 2001; Manji et al., 2012). Acetyl-L-carnitine levels appear to be lower in treatment-resistent MDD patients (n=71) compared to healthy controls (n=45) (Nasca et al., 2018). Moreover, patients undergoing hemodialysis who were administered L-carnitine had increased total, free, and acylcarnitine levels in plasma and an associated decrease in depression rating scales scores (Tashiro et al., 2017). To date, we are unaware of any studies that link acylcarnitine levels or their changes under antidepressant treatment with clinical phenotypes in patients with major depression.

This study addresses the following questions:

1. Are there unique metabolomic patterns that differentiate each pure MDD phenotype (CD+; NVSM+; ANX+) from the others at baseline?
2. Are there unique metabolomic patterns that differentiate each pure phenotype after eight weeks of treatment with an SSRI?
3. Are there unique changes in acylcarnitine profiles over eight weeks of SSRI treatment that differentiate the three phenotypes?

## Materials and Methods

### Study Design and Participants

The sample comprised 240 MDD participants from the Mayo Clinic NIH-Pharmacogenomics Research Network - Antidepressant Medication Pharmacogenomics Study (PGRN-AMPS) (Mrazek et al., 2014). Participant selection, symptomatic evaluation, and blood sample collection for the PGRN-AMPS clinical trial have been described in our previous work (Gupta et al., 2016; Ji et al., 2014; Mrazek et al., 2014; Schiepers, Wichers, & Maes, 2005). MDD symptoms were assessed using the HRSD_17_ at baseline and after eight weeks of treatment with citalopram or escitalopram. Blood samples for metabolomics were collected at these same time points. The data extraction protocol followed the STROBE guidelines (von Elm, Altman, Egger, & et al., 2007).

The study was approved and monitored by the Institutional Review Boards of Mayo Clinic and conformed to the standards of the Declaration of Helsinki. All participants provided written informed consent prior to participation in this study.

### Identifying Subtypes of Major Depression

We previously defined three clinical phenotypes that aligned with RDoC conceptual models (Insel et al., 2010) using the individual HRSD_17_ item scores from participants with MDD (Ahmed et al., 2018). The CD+ phenotype represents the RDoC domain of negative valence systems, and the constructs of loss and reward learning. We defined it based on symptoms’ severity of items #1 (depressed mood) and #7 (work and activities) on the HRSD_17_. The NVSM+ phenotype represents the RDoC domain of arousal and regulatory systems, and the construct of sleep-wakefulness. We defined it based on symptoms’ severity of items #6 (late insomnia) and #12 (somatic symptoms gastrointestinal) on the HRSD_17_. The ANX+ phenotype reflects the RDoC domain of negative valence systems, and the construct of potential threat (“Anxiety”). We defined it based on symptoms’ severity of items #9 (agitation), #10 (psychological anxiety), #11(somatic anxiety), and #15 (hypochondriasis) on the HRSD_17_. **Supplemental Table 1, Figure 1**).

**Figure 1.**
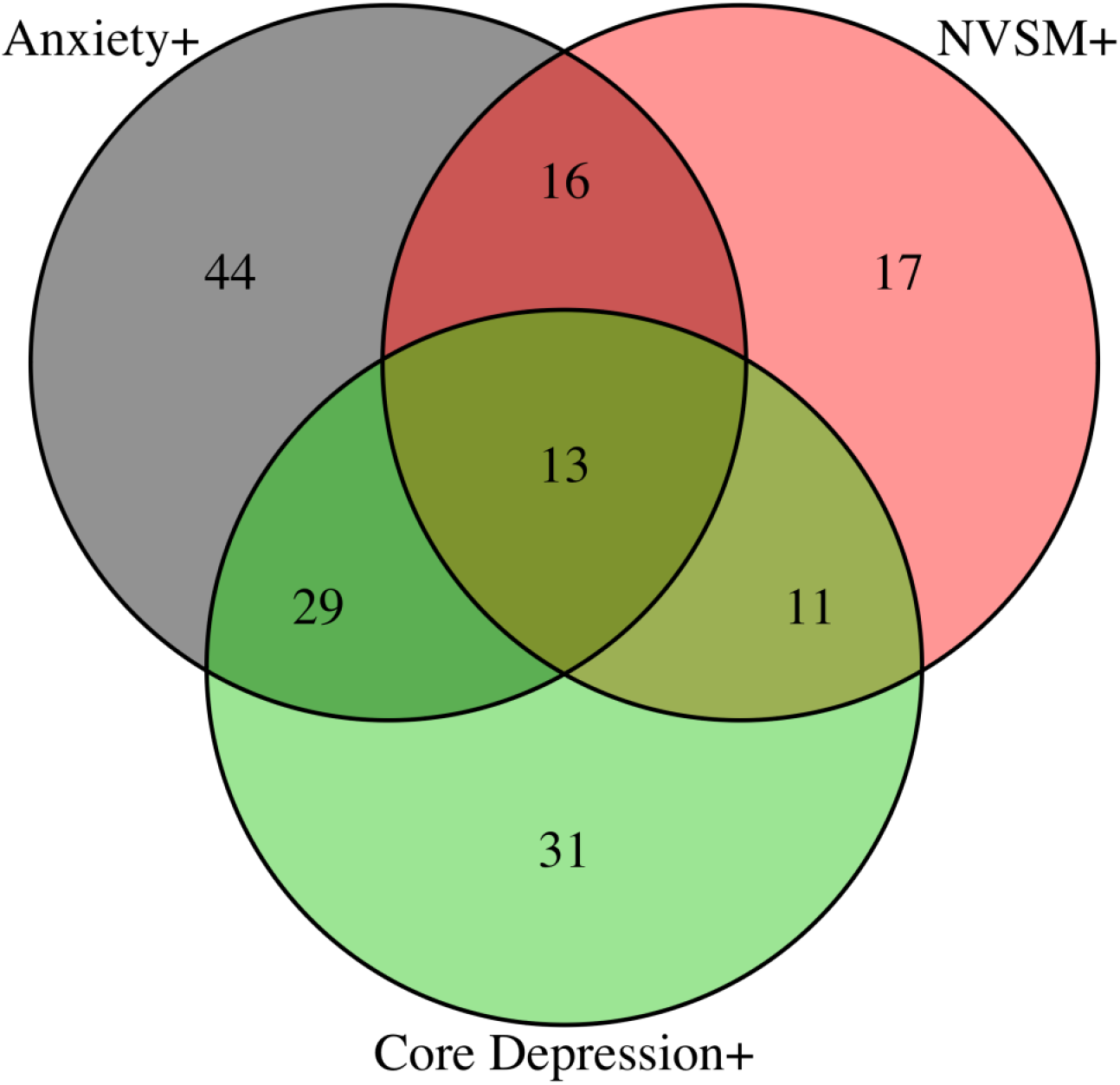
Venn Diagram of the Overlap between Positive MDD Phenotype [Core Depression (CD+), Neurovegetative Symptoms of Melancholia (NVSM+), and Anxiety (ANX+)]. Circles represent the group in each phenotype (CD+, NVSM+, and ANX+). The numbers inside overlapping circles represent participants who met criteria for two, or all three, of the phenotypes. The total number of participants with each pure phenotype was CD+ (n=31), NVSM+ (n=17) and ANX+ (n=44).

### Metabolomic Profiling

Metabolites were measured with a targeted metabolomics approach using the AbsoluteIDQ^®^ p180 Kit (BIOCRATES Life Science AG, Innsbruck, Austria), with an ultra-performance liquid chromatography (UPLC)/MS/MS system [Acquity UPLC (Waters), TQ-S triple quadrupole MS/MS (Waters)], which provides semi-quantitative measurements of up to 23 endogenous acylcarnitines as well as multiple analytes from various other classes including amino acids, biogenic amines, phophatidylcholines, and sphingomyelins.

In this report, we focused solely on acylcarnitines. The AbsoluteIDQ^®^ p180 kit has been fully validated according to European Medicine Agency guidelines. Additionally, the kit plates include an automated technical validation protocol to approve the validity of the run and provide verification of the actual performance of the applied quantitative procedure including instrumental analysis. The technical validation of each analyzed kit plate was performed using MetIDQ^®^ software based on results obtained, and defined acceptance criteria for blank, zero samples, calibration standards and curves, low/medium/high-level quality control (QC) samples, and measured signal intensity of internal standards over the plate. This platform has been used in numerous publications, including several studies of MDD (Moaddel et al., 2018) (Baranyi et al., 2018). De-identified samples were analyzed following the manufacturer’s protocol, with the metabolomics labs blinded to diagnosis and clinical data. The list of acylcarnitines reported in the present study, which does not include several in the p180 kit deemed to be irrelevant to mitochondrial metabolism, is shown in **Supplemental Table 2**.

### Data Analyses

#### Preprocessing

All metabolite data were first checked for missing values (< limit of detection [<LOD]), and metabolites with >40% missing values were excluded from subsequent analysis. Each assay plate included a set of duplicates obtained by combining approximately 10 μl from the first 76 samples in the study (QC pool duplicates) to enable appropriate inter-plate abundance scaling based specifically on this cohort of samples. To adjust for batch effects, a correction factor for each metabolite in a specific plate was obtained by dividing the metabolite’s statistical process quality control (SPQC) global average by the SPQC average within the plate. <LOD values were imputed using each metabolite’s LOD/2 value followed by log2 transformation. We checked for the presence of multivariate outlier samples by evaluating the squared Mahalanobis distance of samples in each platform. Samples with Mahalanobis distances that exceeded the upper 0.05/n (with n: number of samples to adjust for multiple comparisons by Bonferroni correction) critical value of the Chi-squared distribution with m degrees of freedom, in which m is the number of metabolites in each platform, were flagged as outliers. An additional 15 samples were removed after they were determined to be multivariate outliers. This resulted in an analysis data set containing 240 participants, 537 samples and 23 metabolites.

#### Statistical Analysis

Differences in demographic and clinical characteristics, and in HRSD_17_ scores, across the phenotypes were evaluated using the F-test (for continuous variables) and Pearson Chi-squared test (for categorical variables). All analyses were performed in a metabolite-wise (univariate analysis) manner. The presence of each pure phenotype (CD+, NVSM+ and ANX+) for each participant at baseline was stored as three binary variables. Baseline and week eight metabolite concentrations, and changes in metabolite concentrations after SSRI treatment, were tested. To examine the statistical significance of differences in metabolite levels between phenotypes, at baseline and at eight weeks, we fitted linear mixed effect models with participants as the random variable. Log2 metabolite levels were used as the dependent variable. Phenotype (8-level categorical variable: all possible combinations of [CD+|CD−][NVSM+|NVSM−][ANX+|ANX−]) and time of visit (2-level categorical variable: baseline; week eight) were used as independent variables while adjusting for age, sex, HRSD_17_ scores at time of visit, and specific antidepressant (escitalopram or citalopram). We used the “emmeans” R package was employed to compute the least squared means of the contrasts of interest and their corresponding p-values at baseline and at eight weeks (i.e., 1: CD+ vs. NVSM+; 2: CD+ vs. ANX+; 3: NVSM+ vs. ANX+). To examine the statistical significance of temporal metabolite concentration changes over eight weeks across phenotypes, linear mixed effect models (participant as random) were fitted with log2-fold metabolite levels as the dependent variable and time of visit (2-level categorical variable: baseline; week eight) as the independent variable; the models were adjusted for age, sex, baseline HRSD_17_ score, and specific antidepressant (escitalopram or citalopram).

## Results

### Participant Characteristics

Plasma metabolite data were available from 240 MDD participants. At baseline, age, gender, and baseline HRSD_17_ scores did not significantly differ across the three groups. The demographic and clinical characteristics of the study sample are detailed in **Table 1**.

**Table 1:** Baseline Demographics.

| **Feature** | **All Phenotype**<br>**(N=92)** | **CD+**<br>**(N=31)** | **NVSM+**<br>**(N=17)** | **ANX+**<br>**(N=44)** |
| --- | --- | --- | --- | --- |
| AGE Mean (SD) | 39.8 (13.6) | 43.1 (13.9) | 36.6 (14.8) | 36.8 (12.3) |
| Gender Male % (N) | 33% (56) | 32% (10) | 12% (2) | 34% (15) |
| Baseline HRSD<sub>17</sub> Mean (SD) | 20.3 (3.7) | 21.5 (3.9) | 21.5 (3.10) | 22.3 (3.36) |

*Abbreviation:* ANX: Anxious Depression phenotype, CD: Core Depression phenotype, HRSD<sub>17</sub>: The 17-item Hamilton Rating Scale for Depression; NVSM: Neurovegetative Symptoms of Melancholia phenotype

#### 1. At baseline, acylcarnitine metabolomic patterns that differentiate each pure phenotype (CD+; NVSM+; ANX+) from the others

CD+ participants demonstrated significantly lower levels of C0, C3, C5:1, C5-DC/C6-OH, C16-OH, and C18 acylcarnitines compared to ANX+ participants. Moreover, C5-DC/C6-OH and C16-OH acylcarnitines levels were significantly lower, and C10 significantly higher, in the NVSM+ participants compared to the ANX+ participants. **Figure 2** shows the baseline and week eight acylcarnitine metabolite concentrations for each of the three phenotypes. **Supplemental Table 3** shows the baseline and week eight differences between each phenotype and significant p-values.

**Figure 2:**
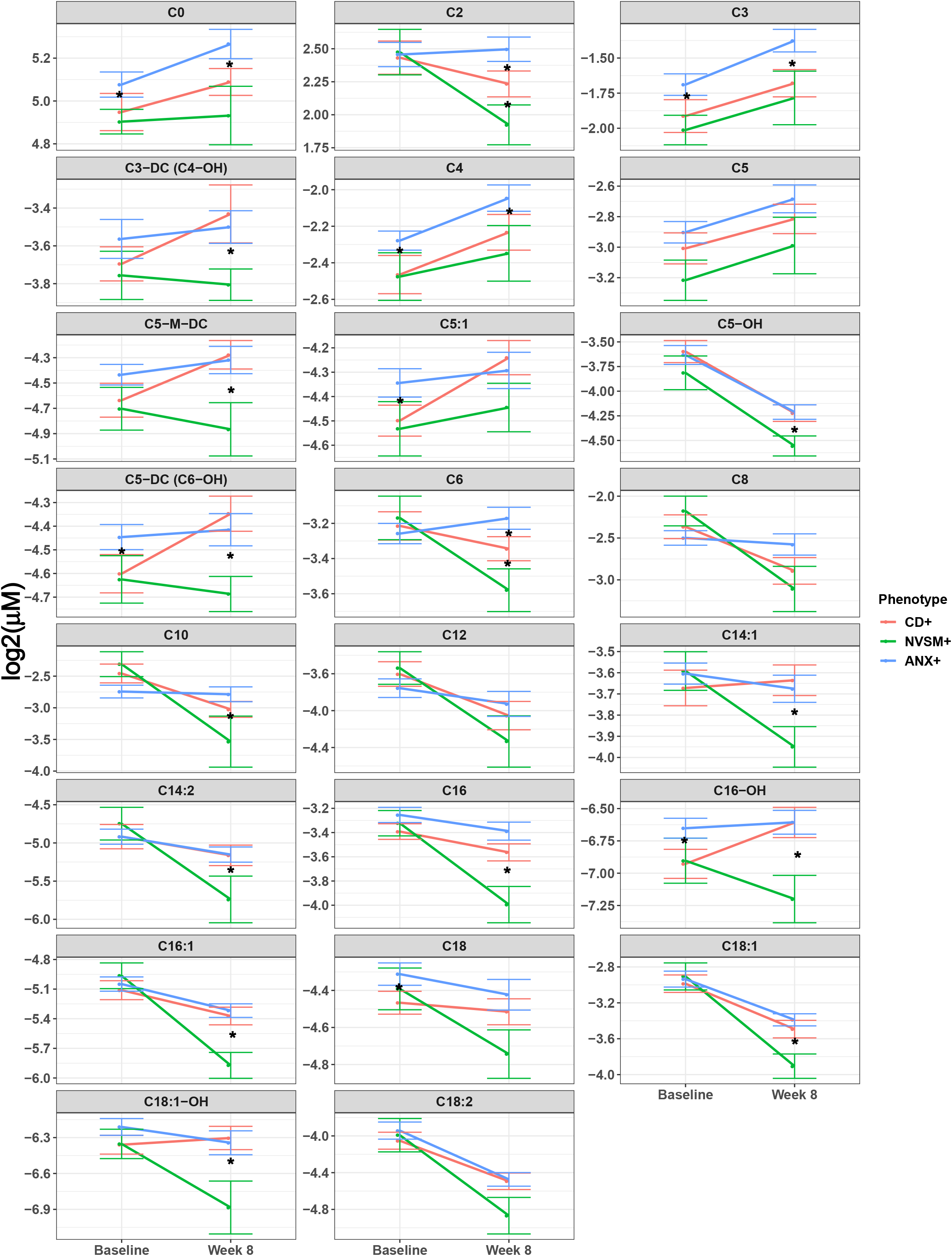
Short-, Medium- & Long-Chain Acylcarnitines — Baseline and 8 Weeks Stratified by Phenotype (+). These plots show the baseline and week-eight acylcarnitine metabolite concentrations for each of the three phenotypes (CD+: Core Depression, ANX+: Anxiety, NVSM+: Neurovegetative Symptom of Melancholia). Asterisks indicate statistical significance of mean differences between the two groups (unadjusted *p*<0.05) at each visit. Error bars represent standard error of the means. P-values were obtained from linear mixed effect models corrected for age, sex, drug and 17-item Hamilton Rating Scale for Depression scores.

#### 2. At eight weeks, acylcarnitine metabolomic patterns that differentiate each pure phenotype (CD+; NVSM+; ANX+) from the others

At eight weeks, CD+ participants had significantly higher levels of C5-DC/C6-OH, C5-M-DC, C6, C14:1, C16, C16-OH, C16:1, and C18:1-OH acylcarnitines compared to NVSM+ participants. CD+ participants had significantly lower levels of C2, C3 and C6 acylcarnitines compared to ANX+ participants. NVSM+ participants had significantly lower levels of C0, C2, C3, C5-DC/C6-OH, C5-M-DC, C6, C10, C14:1, C16, C16-OH, C16:1 and C18:1-OH acylcarnitines compared to ANX+ participants. (**Figure 2, Supplemental Table 3**).

#### 3. Baseline to week eight changes in acylcarnitine metabolomic patterns that differentiate each pure phenotype (CD+; NVSM+; ANX+) from the others

Several short-, medium-, and long-chain acylcarnitines significantly changed in each of the three phenotypes over the eight-week course of SSRI treatment. The heatmap in **Figure 3** illustrates key changes in acylcarnitine evaluated after eight weeks of treatment. Only CD+ participants had significantly increased levels of five short-chain and one long-chain acylcarnitines. Moreover, NVSM+ participants had significantly decreased levels of C2, C6 and six long-chain acylcarnitines (C14:1, C14:2, C16, C18, C18:1, C18:1-OH and C18:2); C5 was significantly increased only in ANX+ participants. C8, C10, and C12 levels significantly decreased in both CD+ and NVSM+ participants, and C0, C3, and C4 levels significantly increased in both CD+ and ANX+ participants. Worth noting, all three phenotypes showed significant decreases in C5-OH, C16:1, C18:1 and C18:2 acylcarnitine levels. Significant p-values ranged from <0.05 to <0.001. **Figure 3 and Supplemental Table 4** show change of metabolite levels from baseline to eight weeks of SSRI treatment among the three phenotypes and significant p-values (rows represent the acylcarnitine and columns represent the phenotype).

**Figure 3.**
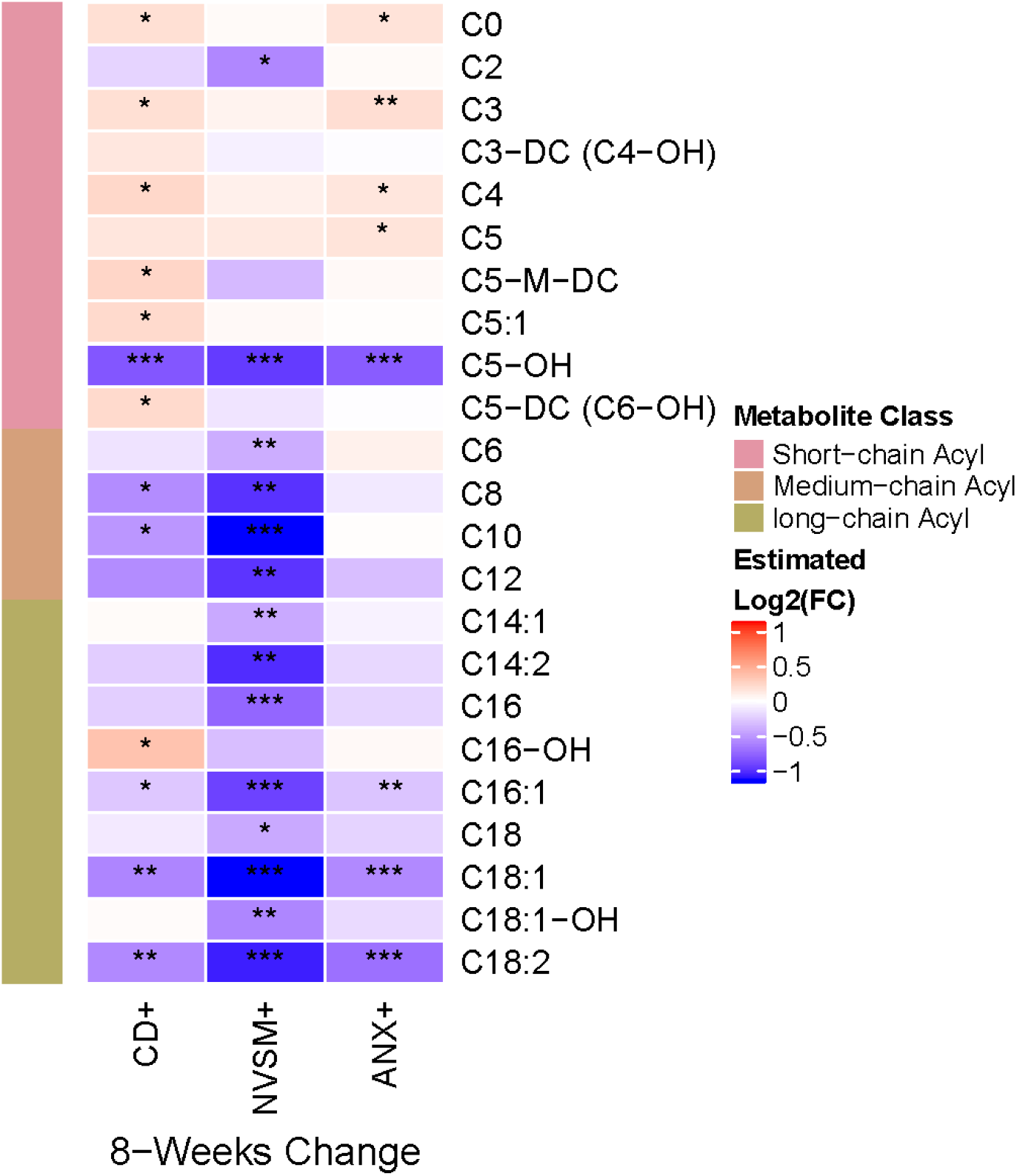
Metabolic Signature of Eight Weeks of Exposure to SSRI. The heatmap depicts the log2 fold change of metabolite levels from baseline to week eight of SSRI treatment. CD+: Core Depression, ANX+: Anxiety, NVSM+: Neurovegetative Symptom of Melancholia. P-values were obtained using linear mixed effect models controlling for age, sex and baseline 17-item Hamilton Rating Scale for Depression scores. Red indicates an increase and blue indicates a decrease in metabolite levels over eight weeks of treatment; *:p-value<0.05, **:p-value<0.01 and ***:p-value<0.001.

## Discussion

This study assessed whether three symptomatically defined phenotypes of MDD could be differentiated based on acylcarnitine profiles at baseline, after eight weeks of citalopram/escitalopram treatment, and temporal changes in the 23 acylcarnitines from baseline to eight weeks of treatment. Both the cross-sectional levels and the changes in acylcarnitine levels distinguished the three pure phenotypes. In the temporal analyses, eight weeks of SSRI treatment was associated with significant increases in short-chain acylcarnitines in the CD+ and ANX+ phenotypes, whereas in the NVSM+ phenotype the treatment was associated with significant decreases in medium- and long-chain acylcarnitines.

After eight weeks of SSRI treatment, acylcarnitine levels in NVSM+ participants were significantly lower than in ANX+ participants. The NVSM+ group showed a decrease in the levels of C2 acylcarnitines after eight weeks of citalopram/escitalopram. The NVSM+ phenotype was defined by loss of sleep and appetite. In a prior study, C2 levels were lower in treatment-resistant depressed patients (n=71) compared to healthy controls (n=45) (Nasca et al., 2018). Another study found a decrease in C2 concentrations after ketamine treatment (n=29) compared to placebo (n=29) (Moaddel et al., 2018). Moreover, increased levels of C2 have been associated with fasting state (Costa, De Almeida, Jakobs, Poll-The, & Duran, 1999; Frohlich, Seccombe, Hahn, Dodek, & Hynie, 1978; Hoppel & Genuth, 1980). Of interest, people with narcolepsy, which is characterized by abnormal sleep, may have low levels of C2 and increased levels of carnitine palmitoyltransferase 1B, which suggests dysfunction in the fatty acid β-oxidation pathway (Miyagawa et al., 2011). Davies and colleagues found increases in this C2 in sleep deprivation (Davies et al., 2014) and they concluded that any changes in the acylcarnitines involved in fatty acid β-oxidation are implicated in sleep regulation (Davies et al., 2014), which is a key component of the NVSM+ phenotype. Our results also show that the NVSM+ and CD+ phenotypes were associated with decreased levels of C8, C10 and C12 after eight weeks of treatment. Furthermore, the changes from baseline to eight weeks included decreases in several medium- and long-chain acylcarnitines in both the NVSM+ and CD+ phenotypes.

The ANX+ phenotype had significantly higher levels of acylcarnitines at baseline and after treatment compared to the CD+ and NVSM+ phenotypes. Short- and long-chain acylcarnitine levels were also significantly different between the CD+ and ANX+ phenotypes. After eight weeks of treatment, short-chain acylcarnitine levels were significantly lower in the CD+ phenotype compared to the ANX+ phenotype. Despite these different symptom profiles, the metabolomic profiles of the CD+ and ANX+ phenotypes overlapped. C0, C3 and C4 levels significantly increased in both phenotypes, although C5 was only increased in the ANX+ phenotype. The CD+ phenotype is defined by patients who are sad and anhedonic, and the ANX+ phenotype is defined by patients who have symptoms of psychological and somatic anxiety and agitation. C0 levels are inversely correlated with self-rated depressive symptoms (Fukami et al., 2014). Tashiro and colleagues have suggested that C4 levels are associated with depressive symptoms (Tashiro et al., 2017). Moreover, hemodialysis patients had a decrease in their self-rated depression scale associated with increased acylcarnitine levels of C4 and C5.

To summarize, the current data demonstrated that three clinically-defined depression phenotypes have distinct patterns of acylcarnitine levels at baseline and after eight weeks of antidepressant treatment. Also, these phenotypes were uniquely and distinctly related to changes in acylcarnitine levels induced by SSRI exposure. These findings and the relationship of acylcarnitine levels with mitochondrial fatty acid β–oxidation and branched-chain amino acid catabolism support efforts toward mapping the global biochemical changes and networks in MDD. Further, these findings may reflect changes in mitochondrial function or ATP production in patients with MDD. Linking acylcarnitine metabolomic data to genetic data has proven to be a powerful approach for highlighting variation in predicting antidepressant response among depressed individuals (Ji et al., 2011; Neavin, Kaddurah-Daouk, & Weinshilboum, 2016). These findings may help to develop a metabolomic profile or ‘metabotype’ of MDD patients, as indexed by an electrochemistry metabolomics platform total output (digital map), with the aim of improving subtype classification of the MDD syndrome.

### Limitations

The study has several limitations. Analyses were performed on a subset of MDD participants, thereby limiting the generalizability of the findings. Some of the reported acylcarnitines, especially dicarboxylic and hydroxylated species (C3DC, C5DC, C5MDC, C16OH, C18OH)— are canonically elevated only in patients with rare inborn errors of metabolism (see Supplemental Table 2), and are of very low concentrations in most individuals. The flow-injection MS/MS method used in this study may lack molecular specificity to confirm exact structure for low-level species, even though the measures reported herein show excellent technical reproducibility. Importantly, although we reported many acylcarnitines which are considered ‘low abundance (i.e. <0.1 μM), our average reported values for each analyte in the cohort are below the pathological clinical reference threshold reported by Mayo Clinic Laboratories (Mayo Clinic Laboratories, 2017). Confirmation of exact molecular speciation of the low-abundance acylcarnitines reported in this study may benefit from utilization of an assay with better molecular specificity, such as a research-grade LC-MS/MS assay reported previously(Minkler et al., 2015). The presence of outlier samples in our cohort might represent subtypes of MDD that will require larger sample sizes to characterize more fully, or may represent other metabolic disorders in those individuals. Our study lacked inclusion of a healthy control group with which to compare and contrast the acylcarnitine profiles of the MDD phenotypes, and allow examination of whether these metabolite changes are due to the drug effect or are unique to the phenotype.

### Conclusions

Baseline and week eight acylcarnitine concentrations, evaluated both cross-sectionally and by temporal change, indicated unique acylcarnitine signatures that distinguish clinically-defined MDD phenotypes from each other. This study supports the possibility of using acylcarnitines as a tool for understanding personal variation in MDD drug response; this moght allow for sub-classifications of MDD to CD+, NVSM+ and ANX+ patients and the linking of acylcarnitine changes to mitochondrial fatty acid β–oxidation and branched-chain amino acid catabolism.

## Acknowledgements

Support for this work was provided by NIMH through the following grants R01MH108348, Dr. Kaddurah Daouk by the National Institute on Aging grant R01AG046171 and the AD Metabolomics Consortium is funded as part of NIA’s national initiatives AMP-AD and M2OVE-AD (R01 AG046171 & U01 AG061359, RF1 AG051550). RO1 MH080880; PI: W Edward Craighead, PhD provided funding to treat non-remitters to the first treatment with combination medication and psychotherapy, to allow follow-up of patients for up to two years to identify predictors of recurrence, and to add patients to the sample to adequately power these studies. Additional support was received from PHS Grant UL1 RR025008 from the Clinical and Translational Science Award program, National Institutes of Health, National Center for Research Resources, PHS Grant M01 RR0039 from the General Clinical Research Center program, and K23 MH086690 (BWD). We thank Lisa Howerton for her administrative support. We also thank the study participants and their families of the Mayo Pharmacogenomics Research Network-Antidepressant Pharmacogenomics Medication Study (PGRN-AMPS) (NIH grants RO1 GM28157, U19 GM61388). We acknowledge the editorial support of Jon Kilner, MS, MA (Pittsburgh, PA).

## Conflict of Interest

Richard Weinshilboum is a co-founder and stockholder in OneOme, LLC, a pharmacogenomic clinical decision support company. Dr. Kaddurah Daouk is an inventor on patents related to metabolomics applications in the study of CNS diseases and holds equity in Metabolon, Inc., a biotechnology company that provides metabolic profiling capabilities. A. John Rush has received: consulting fees from Akili, Brain Resource Inc., Compass Inc., Curbstone Consultant LLC., Emmes Corp., Johnson and Johnson (Janssen), Liva-Nova, Mind Linc., Sunovion; speaking fees from Janssen and Liva-Nova; and royalties from Guilford Press and the University of Texas Southwestern Medical Center, Dallas, TX (for the Inventory of Depressive Symptoms and its derivatives). Dr. Rush is also named co-inventor on two patents: U.S. Patent No. 7,795,033: Methods to Predict the Outcome of Treatment with Antidepressant Medication and U.S. Patent No. 7,906,283: Methods to Identify Patients at Risk of Developing Adverse Events During Treatment with Antidepressant Medication. Drew Neavin’s stipend has been supported in part by NIH T32 GM072474 and the Mayo Graduate School. Ahmed Ahmed’s research was supported by National Institute of General Medical Sciences of the National Institutes of Health under award number T32 GM008685. Matthias Arnold was supported by National Institute on Aging [R01AG057452, RF1AG051550, R01AG046171], National Institute of Mental Health [R01MH108348], and Qatar National Research Fund [NPRP8-061-3-011]. The funders listed above had no role in the design and conduct of the study; collection, management, analysis, and interpretation of the data; preparation, review, or approval of the manuscript; or the decision to submit the manuscript for publication. Dr. Dunlop has received research support from Acadia, Axsome, Janssen, Takeda and he serves as a consultant to Assurex Health and Aptinyx Inc. Dr. Craighead is a board member of Hugarheill ehf, an Icelandic company dedicated to the prevention of depression, and he receives book royalties from John Wiley & Sons; his research is also supported by the Mary and John Brock Foundation and the Fuqua family foundations; he is a consultant to the George West Mental Health Foundation and is a member of the scientific advisory boards of the AIM for Mental Health Foundation and the Anxiety Disorders Association of America. Dr. Bobo’s research has been supported by the NIMH, AHRQ, Mayo Foundation for Medical Education and Research; he has contributed chapters to UpToDate on the treatment of bipolar disorders. Additional support was received from PHS Grant UL1 RR025008 from the Clinical and Translational Science Award program, National Institutes of Health, National Center for Research Resources, PHS Grant M01 RR0039 from the General Clinical Research Center program, and K23 MH086690 (BWD).

## References

A. John Rush, & Hicham M. Ibrahim. (2018). A Clinician’s Perspective on Biomarkers. FOCUS, 16(2), 124–134. doi:10.1176/appi.focus.20170044

Ahmed, A. T., Frye, M. A., Rush, A. J., Biernacka, J. M., Craighead, W. E., McDonald, W. M., … Dunlop, B. W. (2018). Mapping depression rating scale phenotypes onto research domain criteria (RDoC) to inform biological research in mood disorders. J Affect Disord, 238, 1–7. doi:10.1016/j.jad.2018.05.005

Baranyi, A., Meinitzer, A., Rothenhäusler, H.-B., Amouzadeh-Ghadikolai, O., Lewinski, D. V., Breitenecker, R. J., & Herrmann, M. (2018). Metabolomics approach in the investigation of depression biomarkers in pharmacologically induced immune-related depression. PloS one, 13(11), e0208238–e0208238. doi:10.1371/journal.pone.0208238

Beasley, C. L., Pennington, K., Behan, A., Wait, R., Dunn, M. J., & Cotter, D. (2006). Proteomic analysis of the anterior cingulate cortex in the major psychiatric disorders: Evidence for disease-associated changes. Proteomics, 6(11), 3414–3425. doi:10.1002/pmic.200500069

Bresolin, N., Freddo, L., Vergani, L., & Angelini, C. (1982). Carnitine, carnitine acyltransferases, and rat brain function. Experimental Neurology, 78(2), 285–292. doi:https://doi.org/10.1016/0014-4886(82)90047-4

Calabrese, V., Stella, A. M. G., Calvani, M., & Butterfield, D. A. (2006). Acetylcarnitine and cellular stress response: roles in nutritional redox homeostasis and regulation of longevity genes. The Journal of Nutritional Biochemistry, 17(2), 73–88. doi:https://doi.org/10.1016/j.jnutbio.2005.03.027

Carragher, N., Adamson, G., Bunting, B., & McCann, S. (2009). Subtypes of depression in a nationally representative sample. J Affect Disord, 113(1-2), 88–99. doi:10.1016/j.jad.2008.05.015

Cassano, G. B., Benvenuti, A., Miniati, M., Calugi, S., Mula, M., Maggi, L., … Frank, E. (2009). The factor structure of lifetime depressive spectrum in patients with unipolar depression. J Affect Disord, 115(1-2), 87–99. doi:10.1016/j.jad.2008.09.006

Charney, D. S., & Drevets, W. C. (2002). The neurobiological basis of anxiety disorders. .Philadelphia, PA.: Lippincott Williams & Wilkins:.

Chen, S., Wei, C., Gao, P., Kong, H., Jia, Z., Hu, C., … Xu, G. (2014). Effect of Allium macrostemon on a rat model of depression studied by using plasma lipid and acylcarnitine profiles from liquid chromatography/mass spectrometry. J Pharm Biomed Anal, 89, 122–129. doi:10.1016/j.jpba.2013.10.045

Costa, C. C., De Almeida, I. T., Jakobs, C., Poll-The, B.-T., & Duran, M. (1999). Dynamic changes of plasma acylcarnitine levels induced by fasting and sunflower oil challenge test in children. Pediatric research, 46(4), 440.

Davies, S. K., Ang, J. E., Revell, V. L., Holmes, B., Mann, A., Robertson, F. P., … Skene, D. J. (2014). Effect of sleep deprivation on the human metabolome. Proc Natl Acad Sci U S A, 111(29), 10761–10766. doi:10.1073/pnas.1402663111

Fattal, O., Link, J., Quinn, K., Cohen, B. H., & Franco, K. (2007). Psychiatric comorbidity in 36 adults with mitochondrial cytopathies. CNS Spectr, 12(6), 429–438.

Fritz, I. B., & McEwen, B. (1959). Effects of Carnitine on Fatty-Acid Oxidation by Muscle. Science, 129(3345), 334–335. doi:10.1126/science.129.3345.334

Frohlich, J., Seccombe, D. W., Hahn, P., Dodek, P., & Hynie, I. (1978). Effect of fasting on free and esterified carnitine levels in human serum and urine: correlation with serum levels of free fatty acids and beta-hydroxybutyrate. Metabolism, 27(5), 555–561.

Fukami, K., Yamagishi, S., Sakai, K., Kaida, Y., Minami, A., Nakayama, Y., … Okuda, S. (2014). Carnitine deficiency is associated with late-onset hypogonadism and depression in uremic men with hemodialysis. Aging Male, 17(4), 238–242. doi:10.3109/13685538.2014.888053

Gardner, A., Johansson, A., Wibom, R., Nennesmo, I., von Dobeln, U., Hagenfeldt, L., & Hallstrom, T. (2003). Alterations of mitochondrial function and correlations with personality traits in selected major depressive disorder patients. J Affect Disord, 76(1-3), 55–68.

Gupta, M., Neavin, D., Liu, D., Biernacka, J., Hall-Flavin, D., Bobo, W. V., … Weinshilboum, R. M. (2016). TSPAN5, ERICH3 and selective serotonin reuptake inhibitors in major depressive disorder: pharmacometabolomics-informed pharmacogenomics. Mol Psychiatry, 21(12), 1717–1725. doi:10.1038/mp.2016.6

Harald, B., & Gordon, P. (2012). Meta-review of depressive subtyping models. J Affect Disord, 139(2), 126–140. doi:10.1016/j.jad.2011.07.015

Hoppel, C. L., & Genuth, S. M. (1980). Carnitine metabolism in normal-weight and obese human subjects during fasting. American Journal of Physiology-Endocrinology and Metabolism, 238(5), E409–E415.

Insel, T., Cuthbert, B., Garvey, M., Heinssen, R., Pine, D. S., Quinn, K., … Wang, P. (2010). Research domain criteria (RDoC): toward a new classification framework for research on mental disorders. Am J Psychiatry, 167(7), 748–751. doi:10.1176/appi.ajp.2010.09091379

Ji, Y., Hebbring, S., Zhu, H., Jenkins, G. D., Biernacka, J., Snyder, K., … Weinshilboum, R. M. (2011). Glycine and a glycine dehydrogenase (GLDC) SNP as citalopram/escitalopram response biomarkers in depression: pharmacometabolomics-informed pharmacogenomics. Clin Pharmacol Ther, 89(1), 97–104. doi:10.1038/clpt.2010.250

Ji, Y., Schaid, D. J., Desta, Z., Kubo, M., Batzler, A. J., Snyder, K., … Weinshilboum, R. M. (2014). Citalopram and escitalopram plasma drug and metabolite concentrations: genome-wide associations. Br J Clin Pharmacol, 78(2), 373–383. doi:10.1111/bcp.12348

Jones, L. L., McDonald, D. A., & Borum, P. R. (2010). Acylcarnitines: role in brain. Prog Lipid Res, 49(1), 61–75. doi:10.1016/j.plipres.2009.08.004

Kato, T. (2001). The other, forgotten genome: mitochondrial DNA and mental disorders. Mol Psychiatry, 6(6), 625–633. doi:10.1038/sj.mp.4000926

Manji, H., Kato, T., Di Prospero, N. A., Ness, S., Beal, M. F., Krams, M., & Chen, G. (2012). Impaired mitochondrial function in psychiatric disorders. Nat Rev Neurosci, 13(5), 293–307. doi:10.1038/nrn3229

Mayo Clinic Laboratories. (2017). Acylcarnitines, Quantitative, Plasma. Retrieved from https://www.mayocliniclabs.com/test-catalog/Clinical+and+lnterpretive/82413

Minkler, P. E., Stoll, M. S., Ingalls, S. T., Kerner, J., & Hoppel, C. L. (2015). Quantitative acylcarnitine determination by UHPLC-MS/MS--Going beyond tandem MS acylcarnitine “profiles”. Mol Genet Metab, 116(4), 231–241. doi:10.1016/j.ymgme.2015.10.002

Miyagawa, T., Miyadera, H., Tanaka, S., Kawashima, M., Shimada, M., Honda, Y., … Honda, M. (2011). Abnormally low serum acylcarnitine levels in narcolepsy patients. Sleep, 34(3), 349–353a.

Moaddel, R., Shardell, M., Khadeer, M., Lovett, J., Kadriu, B., Ravichandran, S., … Zarate, C. A. (2018). Plasma metabolomic profiling of a ketamine and placebo crossover trial of major depressive disorder and healthy control subjects. Psychopharmacology (Berl), 235(10), 3017–3030. doi:10.1007/s00213-018-4992-7

Mrazek, D. A., Biernacka, J. M., McAlpine, D. E., Benitez, J., Karpyak, V. M., Williams, M. D., … Katzelnick, D. J. (2014). Treatment outcomes of depression: the pharmacogenomic research network antidepressant medication pharmacogenomic study. J Clin Psychopharmacol, 34(3), 313–317. doi:10.1097/jcp.0000000000000099

Nałęcz, K. A., Miecz, D., Berezowski, V., & Cecchelli, R. (2004). Carnitine: transport and physiological functions in the brain. Molecular Aspects of Medicine, 25(5), 551–567. doi:https://doi.org/10.1016/j.mam.2004.06.001

Nasca, C., Bigio, B., Lee, F. S., Young, S. P., Kautz, M. M., Albright, A., … Rasgon, N. (2018). Acetyl-l-carnitine deficiency in patients with major depressive disorder. Proceedings of the National Academy of Sciences, 115(34), 8627–8632. doi:10.1073/pnas.1801609115

Neavin, D., Kaddurah-Daouk, R., & Weinshilboum, R. (2016). Pharmacometabolomics informs Pharmacogenomics. Metabolomics, 12(7). doi:10.1007/s11306-016-1066-x

Pettegrew, J. W., Levine, J., & McClure, R. J. (2000). Acetyl-L-carnitine physical-chemical, metabolic, and therapeutic properties: relevance for its mode of action in Alzheimer&#39;s disease and geriatric depression. Mol Psychiatry, 5, 616. doi:10.1038/sj.mp.4000805

Rush, A. J., & Ibrahim, H. M. (2018). Speculations on the Future of Psychiatric Diagnosis. J Nerv Ment Dis, 206(6), 481–487. doi:10.1097/nmd.0000000000000821

Schiepers, O. J. G., Wichers, M. C., & Maes, M. (2005). Cytokines and major depression. Progress in Neuro-Psychopharmacology and Biological Psychiatry, 29(2), 201–217. doi:https://doi.org/10.1016/j.pnpbp.2004.11.003

Shug, A. L., Schmidt, M. J., Golden, G. T., & Fariello, R. G. (1982). The distribution and role of carnitine in the mammalian brain. Life Sciences, 31(25), 2869–2874. doi:https://doi.org/10.1016/0024-3205(82)90677-4

Suomalainen, A., Majander, A., Haltia, M., Somer, H., Lonnqvist, J., Savontaus, M. L., & Peltonen, L. (1992). Multiple deletions of mitochondrial DNA in several tissues of a patient with severe retarded depression and familial progressive external ophthalmoplegia. J Clin Invest, 90(1), 61–66. doi:10.1172/jci115856

Tashiro, K., Kaida, Y., Yamagishi, S.-I., Tanaka, H., Yokoro, M., Yano, J., … Fukami, K. (2017). L-Carnitine Supplementation Improves Self-Rating Depression Scale Scores in Uremic Male Patients Undergoing Hemodialysis. Letters in drug design & discovery, 14(6), 737–742. doi:10.2174/1570180814666170216102632

Virmani, A., & Binienda, Z. (2004). Role of carnitine esters in brain neuropathology. Molecular Aspects of Medicine, 25(5), 533–549. doi:https://doi.org/10.1016/j.mam.2004.06.003

von Elm, E., Altman, D. G., Egger, M., & et al. (2007). The strengthening the reporting of observational studies in epidemiology (strobe) statement: Guidelines for reporting observational studies. Annals of Internal Medicine, 147(8), 573–577. doi:10.7326/0003-4819-147-8-200710160-00010

Zanelli, S. A., Solenski, N. J., Rosenthal, R. E., & Fiskum, G. (2005). Mechanisms of Ischemic Neuroprotection by Acetyl-l-carnitine. Annals of the New York Academy of Sciences, 1053(1), 153–161. doi:doi:10.1111/j.1749-6632.2005.tb00021.x

