## Supplemental table 1 for "Acylcarnitine Metabolomic Profiles Inform Clinically-Defined Major Depressive Phenotypes"

**Supplemental Table 1: Phenotype Symptom Constructs and Criteria Thresholds**

| **RDoC Construct** | **Phenotype (Mayo & Emory)** | | **HRSD_17_ items** |
| --- | --- | --- | --- |
| Loss | Core Depression (CD+) | Symptoms | depressed mood #1  work and activities #7 |
|  |  | Criteria definition | score of 3 or 4 on both items #1 and #7 |
| Potential Threat ("Anxiety") | Anxiety (ANX+)* | Symptoms | agitation #9  anxiety psychological #10  anxiety somatic #11  hypochondriasis #15 |
|  |  | Criteria definition | total score ≥6 from items #9, #10, #11, and #15 |
| Sleep-Wakefulness | Neurovegetative Symptoms of Melancholia (NVSM+) | Symptoms | late insomnia #6  somatic gastrointestinal #12 |
|  |  | Criteria definition | score of 1 or 2 on both items #6 and #12 |

*Abbreviations*: HRSD_17_: 17-item Hamilton Rating Scale for Depression; RDoC: Research Domain Criteria
