## Supplemental table 2 for "Acylcarnitine Metabolomic Profiles Inform Clinically-Defined Major Depressive Phenotypes"

**Supplemental Table 2: List of Acylcarnitines Included in the Analysis**

| **Short Name** | **Biochemical Name** | **Source** |
| --- | --- | --- |
| C0 | L-Carnitine | Diet, Endogenous Synthesis |
| C2 | Acetyl-L-carnitine | Multiple |
| C3 | Propionyl-L-carnitine | VAL, ILE, THR, MET |
| C3-DC / C4-OH | (Malonyl-L-carnitine) / Hydroxybutyryl-L-carnitine | Ketones, Rare IEM |
| C4 | Butyryl/Isobutyryl-L-carnitine | FAO, VAL |
| C6 | Hexanoyl-L-carnitine | FAO |
| C5 | Isovaleryl/methylbutyryl-L-carnitine | LEU/ILE |
| C5:1 | Tiglyl/methylcrotyonyl-L-carnitine | LEU/ILE |
| C5-DC / C6-OH | Glutaryl-L-carnitine / Hydroxyhexanoyl-L-carnitine | Rare IEM |
| C5-M-DC | Methylglutaryl-L-carnitine | LEU |
| C5-OH | Hydroxyisovaleryl/methyl-hydroxybutyryl-L-carnitine | LEU/ILE |
| C8 | Octanoyl-L-carnitine | FAO |
| C10 | Decanoyl-L-carnitine | FAO |
| C12 | Dodecanoyl-L-carnitine | FAO |
| C14 | Tetradecanoyl-L-carnitine | FAO |
| C14:1 | Tetradecenoyl-L-carnitine | FAO |
| C14:2 | Tetradecadienyl-L-carnitine | FAO |
| C16 | Hexadecanoyl-L-carnitine | Diet |
| C16:1 | Hexadecenoyl-L-carnitine | Diet |
| C16-OH | Hydroxyhexadecanoyl-L-carnitine | Rare IEM |
| C18 | Octadecanoyl-L-carnitine | Diet |
| C18:1 | Octadecenoyl-L-carnitine | Diet |
| C18:1-OH | Hydroxyoctadecenoyl-L-carnitine | Rare IEM |
| C18:2 | Octadecadienyl-L-carnitine | Diet |

*Abbreviations*: FAO: fatty acid beta-oxidation; IEM: Inborn errors of metabolism; ILE: isoleucine; LEU: leucine; MET: methionine; THR: threonine; VAL: valine
