## Supplemental table 3 for "Acylcarnitine Metabolomic Profiles Inform Clinically-Defined Major Depressive Phenotypes"

**Supplemental Table 3: Significantly Different Metabolite Levels between Phenotype (+) and Each Other at baseline and 8 Weeks**

| **Acylcarnitines** | **Baseline** | | | | | | | | | **8 weeks** | | | | | | | | |
| --- | --- | --- | --- | --- | --- | --- | --- | --- | --- | --- | --- | --- | --- | --- | --- | --- | --- | --- |
|  | **CD+^a^ vs NVSM+^a^** | | | **CD+^a^ vs ANX+^a^** | | | **NVSM+^a^ vs ANX+^a^** | | | **CD+^a^ vs NVSM+^a^** | | | **CD+^a^ vs ANX+^a^** | | | **NVSM+^a^ vs ANX+^a^** | | |
|  | **Estimated Log_2_ Difference** | **SE** | ***p*-value** | **Estimated Log_2_ Difference** | **SE** | ***p*-value** | **Estimated Log_2_ Difference** | **SE** | ***p*-value** | **Estimated Log_2_ Difference** | **SE** | ***p*-value** | **Estimated Log_2_ Difference** | **SE** | ***p*-value** | **Estimated Log_2_ Difference** | **SE** | ***p*-value** |
| **C0** | **-0.05** | **0.10** | **0.636** | **-0.16** | **0.08** | **0.034*** | **-0.12** | **0.09** | **0.214** | *0.09* | *0.12* | *0.435* | **-0.15** | **0.09** | **0.101** | **-0.24** | **0.11** | **0.037***** |
| **C2** | **-0.11** | **0.16** | **0.496** | **-0.10** | **0.13** | **0.411** | *0.01* | *0.15* | *0.966* | *0.22* | *0.20* | *0.268* | **-0.31** | **0.15** | **0.040***** | **-0.54** | **0.20** | **0.006****** |
| **C3** | **-0.01** | **0.14** | **0.947** | **-0.26** | **0.11** | **0.016****** | **-0.25** | **0.13** | **0.055** | *0.09* | *0.17* | *0.584* | **-0.27** | **0.13** | **0.036***** | **-0.36** | **0.17** | **0.028***** |
| **C3-DC/C4-OH** | *0.08* | *0.15* | *0.617* | **-0.13** | **0.12** | **0.264** | **-0.21** | **0.14** | **0.148** | *0.27* | *0.19* | *0.159* | *0.00* | *0.14* | *0.980* | **-0.26** | **0.18** | **0.152** |
| **C4** | **-0.06** | **0.14** | **0.680** | **-0.23** | **0.11** | **0.039***** | **-0.17** | **0.13** | **0.207** | *0.07* | *0.17* | *0.676* | **-0.15** | **0.13** | **0.223** | **-0.22** | **0.16** | **0.165** |
| **C5** | *0.12* | *0.14* | *0.382* | **-0.12** | **0.11** | **0.270** | **-0.24** | **0.13** | **0.068** | *0.14* | *0.17* | *0.424* | **-0.13** | **0.13** | **0.308** | **-0.27** | **0.17** | **0.106** |
| **C5DC/C6-OH** | *0.01* | *0.11* | *0.926* | **-0.20** | **0.08** | **0.021***** | **-0.21** | **0.10** | **0.045***** | *0.32* | *0.13* | *0.016*** | *0.01* | *0.10* | *0.919* | **-0.31** | **0.13** | **0.016****** |
| **C5-M-DC** | *0.09* | *0.20* | *0.662* | **-0.24** | **0.16** | **0.129** | **-0.33** | **0.19** | **0.088** | *0.60* | *0.25* | *0.016*** | **-0.06** | **0.19** | **0.756** | **-0.66** | **0.24** | **0.006****** |
| **C5-OH** | *0.17* | *0.19* | *0.365* | **-0.02** | **0.14** | **0.893** | **-0.19** | **0.18** | **0.285** | *0.28* | *0.24* | *0.231* | **-0.04** | **0.18** | **0.813** | **-0.32** | **0.23** | **0.156** |
| **C5:1** | *0.02* | *0.11* | *0.838* | **-0.17** | **0.09** | **0.044***** | **-0.20** | **0.10** | **0.062** | *0.19* | *0.14* | *0.170* | *0.01* | *0.10* | *0.946* | **-0.18** | **0.13** | **0.173** |
| **C6** | **-0.07** | **0.11** | **0.484** | **-0.03** | **0.08** | **0.717** | *0.04* | *0.10* | *0.658* | *0.16* | *0.13* | *0.221* | **-0.22** | **0.10** | **0.025***** | **-0.38** | **0.13** | **0.003****** |
| **C8** | **-0.27** | **0.21** | **0.216** | *0.03* | *0.17* | *0.840* | *0.30* | *0.20* | *0.140* | *0.11* | *0.27* | *0.674* | **-0.37** | **0.20** | **0.067** | **-0.48** | **0.26** | **0.064** |
| **C10** | **-0.23** | **0.21** | **0.273** | *0.19* | *0.17* | *0.245* | *0.43* | *0.20* | *0.035* | *0.41* | *0.27* | *0.127* | **-0.27** | **0.20** | **0.178** | **-0.68** | **0.26** | **0.009****** |
| **C12** | **-0.14** | **0.22** | **0.539** | *0.07* | *0.17* | *0.702* | *0.20* | *0.21* | *0.335* | *0.23* | *0.28* | *0.403* | **-0.15** | **0.21** | **0.464** | **-0.39** | **0.27** | **0.153** |
| **C14:1** | **-0.08** | **0.11** | **0.450** | **-0.09** | **0.08** | **0.278** | **-0.01** | **0.10** | **0.925** | *0.27* | *0.14* | *0.048** | **-0.01** | **0.10** | **0.892** | **-0.28** | **0.13** | **0.032***** |
| **C14:2** | **-0.21** | **0.22** | **0.348** | **-0.05** | **0.17** | **0.765** | *0.15* | *0.21* | *0.456* | *0.48* | *0.27* | *0.080* | **-0.10** | **0.21** | **0.641** | **-0.58** | **0.27** | **0.031***** |
| **C16** | **-0.12** | **0.13** | **0.340** | **-0.18** | **0.10** | **0.073** | **-0.06** | **0.12** | **0.640** | *0.34* | *0.16* | *0.033** | **-0.20** | **0.12** | **0.089** | **-0.54** | **0.15** | **0.000******* |
| **C16-OH** | **-0.01** | **0.16** | **0.948** | **-0.31** | **0.12** | **0.013****** | **-0.30** | **0.15** | **0.047***** | *0.58* | *0.20* | *0.004*** | **-0.03** | **0.15** | **0.842** | **-0.61** | **0.19** | **0.002****** |
| **C16:1** | **-0.17** | **0.14** | **0.231** | **-0.13** | **0.11** | **0.244** | 0.04 | 0.14 | 0.758 | *0.40* | *0.18* | *0.026** | **-0.11** | **0.14** | **0.399** | **-0.52** | **0.17** | **0.003****** |
| **C18** | **-0.15** | **0.12** | **0.219** | **-0.20** | **0.09** | **0.039***** | **-0.05** | **0.12** | **0.688** | *0.13* | *0.15* | *0.386* | **-0.10** | **0.11** | **0.365** | **-0.23** | **0.14** | **0.110** |
| **C18:1** | **-0.18** | **0.18** | **0.306** | **-0.13** | **0.14** | **0.332** | *0.05* | *0.17* | *0.775* | *0.30* | *0.22* | *0.180* | **-0.15** | **0.17** | **0.372** | **-0.45** | **0.22** | **0.038*** |
| **C18:1-OH** | **-0.02** | **0.14** | **0.876** | **-0.19** | **0.11** | **0.074** | **-0.17** | **0.13** | **0.191** | *0.52* | *0.17* | *0.003*** | **-0.02** | **0.13** | **0.897** | **-0.54** | **0.17** | **0.001******* |
| **C18:2** | **-0.14** | **0.19** | **0.452** | **-0.17** | **0.15** | **0.266** | **-0.02** | **0.18** | **0.905** | *0.29* | *0.24* | *0.239* | **-0.06** | **0.18** | **0.756** | **-0.34** | **0.23** | **0.146** |

The increase or decrease in metabolites concentration is shown by italics or bold fonts; Italics: if the 1^st^ Phenotype+ in column is larger than 2^nd^ Phenotype+; Bold: if the 1^st^ Phenotype+ in column is smaller than 2^nd^ Phenotype+; **^a^**CD: Core depression, ANX: Anxiety, NVSM: Neurovegetative Symptom of Melancholia; Linear mixed effect models were used to assess the significance of visit by treatment outcome interaction. Models were adjusted for age, sex, treatment and baseline 17-item Hamilton Rating Scale for Depression score. Significance level were set to ɑ=0.05 not corrected for Bonferroni corrections; *p <0.05;** p ≤0.01; ***p ≤0.001 and highlighted with grey.
