## Supplemental table 4 for "Acylcarnitine Metabolomic Profiles Inform Clinically-Defined Major Depressive Phenotypes"

**Supplemental Table 4: Baseline to Eight Weeks Metabolite Levels Changes among Phenotype (+)**

| **Acylcarnitines** | **Baseline** | | | | | | | | |
| --- | --- | --- | --- | --- | --- | --- | --- | --- | --- |
|  | **CD+^a^** | | | **NVSM+^a^** | | | **ANX+^a^** | | |
|  | **Estimated Log_2_ Difference** | **SE** | ***p*-value** | **Estimated Log_2_ Difference** | **SE** | ***p*-value** | **Estimated Log_2_ Difference** | **SE** | ***p*-value** |
| **C0** | 0.14 | 0.07 | 0.046* | -0.01 | 0.10 | 0.956 | 0.12 | 0.06 | 0.040* |
| **C2** | -0.19 | 0.13 | 0.142 | -0.54 | 0.19 | 0.004** | 0.01 | 0.11 | 0.901 |
| **C3** | 0.24 | 0.11 | 0.027* | 0.16 | 0.15 | 0.296 | 0.25 | 0.09 | 0.007** |
| **C3-DC/C4-OH** | 0.17 | 0.12 | 0.165 | 0.01 | 0.17 | 0.959 | 0.04 | 0.10 | 0.721 |
| **C4** | 0.23 | 0.09 | 0.012** | 0.11 | 0.13 | 0.380 | 0.16 | 0.08 | 0.043* |
| **C5** | 0.19 | 0.11 | 0.093 | 0.20 | 0.16 | 0.204 | 0.20 | 0.10 | 0.033* |
| **C5DC/C6-OH** | 0.22 | 0.08 | 0.007** | -0.07 | 0.12 | 0.534 | 0.02 | 0.07 | 0.795 |
| **C5-M-DC** | 0.31 | 0.16 | 0.047* | -0.16 | 0.22 | 0.483 | 0.14 | 0.13 | 0.309 |
| **C5-OH** | -0.60 | 0.17 | 0.000*** | -0.70 | 0.24 | 0.004** | -0.58 | 0.15 | 0.000*** |
| **C5:1** | 0.22 | 0.09 | 0.012** | 0.07 | 0.12 | 0.569 | 0.04 | 0.07 | 0.593 |
| **C6** | -0.12 | 0.09 | 0.185 | -0.35 | 0.12 | 0.004** | 0.08 | 0.07 | 0.316 |
| **C8** | -0.50 | 0.18 | 0.006** | -0.88 | 0.26 | 0.001*** | -0.10 | 0.16 | 0.532 |
| **C10** | -0.51 | 0.17 | 0.004** | -1.17 | 0.25 | 0.000*** | -0.05 | 0.15 | 0.757 |
| **C12** | -0.42 | 0.19 | 0.026* | -0.77 | 0.27 | 0.005** | -0.20 | 0.16 | 0.224 |
| **C14:1** | 0.02 | 0.09 | 0.854 | -0.34 | 0.13 | 0.009** | -0.06 | 0.08 | 0.426 |
| **C14:2** | -0.24 | 0.18 | 0.184 | -0.95 | 0.26 | 0.000*** | -0.20 | 0.16 | 0.202 |
| **C16** | -0.16 | 0.11 | 0.126 | -0.61 | 0.15 | 0.000*** | -0.13 | 0.09 | 0.139 |
| **C16-OH** | 0.32 | 0.13 | 0.017** | -0.26 | 0.19 | 0.170 | 0.04 | 0.11 | 0.742 |
| **C16:1** | -0.25 | 0.12 | 0.040* | -0.84 | 0.18 | 0.000*** | -0.27 | 0.11 | 0.012** |
| **C18** | -0.04 | 0.09 | 0.701 | -0.30 | 0.13 | 0.025* | -0.13 | 0.08 | 0.112 |
| **C18:1** | -0.49 | 0.16 | 0.002** | -0.97 | 0.23 | 0.000*** | -0.47 | 0.14 | 0.001*** |
| **C18:1-OH** | 0.04 | 0.12 | 0.744 | -0.49 | 0.17 | 0.003** | -0.14 | 0.10 | 0.176 |
| **C18:2** | -0.42 | 0.17 | 0.014** | -0.85 | 0.24 | 0.001*** | -0.53 | 0.15 | 0.000*** |

**^a^**CD: Core depression, ANX: Anxiety, NVSM: Neurovegetative Symptom of Melancholia; log2 fold change of metabolite levels from baseline to eight weeks of SSRI treatment. P-values were obtained using linear mixed effect models controlling for age, sex, treatment and baseline 17-item Hamilton Rating Scale for Depression score. Significance level were set to ɑ=0.05 not corrected for Bonferroni corrections; *p <0.05;** p ≤0.01; ***p ≤0.001 and highlighted with grey.
